# Mining human prostate cancer datasets: The “camcAPP” shiny app

**DOI:** 10.1101/090589

**Authors:** Mark J Dunning, Sarah L Vowler, Emilie Lalonde, Helen Ross-Adams, Ian G Mills, Andy G Lynch, Alastair D Lamb

## Abstract

Obtaining access to robust, well-annotated human genomic datasets is an important step in demonstrating relevance of experimental findings and, often, in generating the hypotheses that led to those experiments being conducted in the first place. We recently published data from two cohorts of men with prostate cancer who had undergone prostatectomy, men from Cambridge, UK and Stockholm, Sweden.^1^ We considered how we might best share our output with those who wish to interrogate the data with their own ideas, gene lists and clinical questions. We recognised that finding, down-loading, processing and assimilating any such dataset into a usable format is daunting and may put off even the more persistent researcher. We also felt that interrogation tools generated to date (e.g. cBioPortal) lack functionality as they either cover too many organ types, or fall short in terms of clinical annotation. We therefore determined to produce an accessible web-based platform that would permit straightforward interrogation of these datasets with individual gene identifiers or gene sets. Furthermore, we decided to include additional publicly accessible human prostate cancer sets in order to increase the number of samples available and provide a degree of validation of any observations made across independent cohorts. We included a number of prominent publicly available sets with both gene expression and copy number data leading to a cohort of almost 500 men.^1-3^ We also included a small landmark series of expression data.^4^ These studies are summarised in **Table 1**. We plan to include additional studies in the app as well-annotated datasets become publicly available.

**Table 1.**
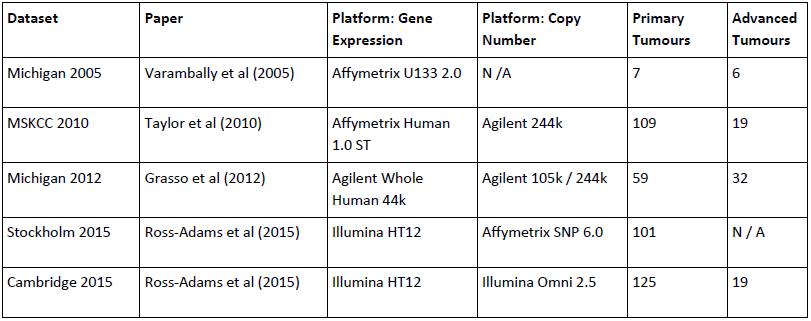
Summary of studies included in the camcAPP at initial release. Primary Tumours = tissue taken from radical prostatectomy specimens in men with confirmed organ-confined disease. Advanced Tumours = tissue from channel transurethral resection of the prostate (chTURP) or prostatectomy in men with metastatic disease.

| Dataset | Paper | Platform: Gene Expression | Platform: Copy Number | Primary Tumours | Advanced Tumours |
| --- | --- | --- | --- | --- | --- |
| Michigan 2005 | Varambally et al (2005) | Affymetrix U133 2.0 | N / A | 7 | 6 |
| MSKCC 2010 | Taylor et al (2010) | Affymetrix Human 1.0 ST | Agilent 244k | 109 | 19 |
| Michigan 2012 | Grasso et al (2012) | Agilent Whole Human 44k | Agilent 105k / 244k | 59 | 32 |
| Stockholm 2015 | Ross-Adams et al (2015) | Illumina HT12 | Affymetrix SNP 6.0 | 101 | N / A |
| Cambridge 2015 | Ross-Adams et al (2015) | Illumina HT12 | Illumina Omni 2.5 | 125 | 19 |

An important finding in our recent study was that prostate cancer could be divided into five distinct molecular subgroups based on stratification by a small number of copy number features which were also associated with expression change. These groups had different clinical outcomes. We wanted the app to allow researchers to determine the mean expression level or copy number status of a single gene or geneset in prostates from men divided either according to clinical categories (Gleason score, biochemical relapse status or tumour type) or according to molecular subgroups. These subgroups could either be pre-defined molecular groups published in the relevant papers, or *de novo* subgroups generated by hierarchical clustering based on an uploaded geneset.

We searched for other tools that are already available for this purpose. Although no such site exists for assessment of subgroup patterns or combined expression and copy number profiles, the Memorial Sloane Kettering Cancer Centre (MSKCC) and Michigan data (**Table 1**) can be analysed as part of cBioPortal^5^ along with the recently published prostate TCGA dataset.^6^

Here we introduce the camcAPP. This is implemented as a Shiny^7^ application in R^8^. Shiny allows the non-specialist bioinformatician to create publication-ready figures and tables through an intuitive interface to the underlying R code. The source code for the entire app is also available through github^9^ (https://github.com/crukci-bioinformatics/camcAPP). The dplyr^10^ package is used throughout for efficient data manipulation, and graphics are generated using ggplot2^11^. The datasets themselves are available via Gene Expression Omnibus. Using the GEOquery^12^ package, we downloaded the datasets and converted them into a format compatible with the Bioconductor^13^ project. The resulting packages are available as experimental data packages in Bioconductor.

After selecting a dataset of interest, and uploading a list of genes (**Figure 1**), the following analyses can be performed:-

**Figure 1.**
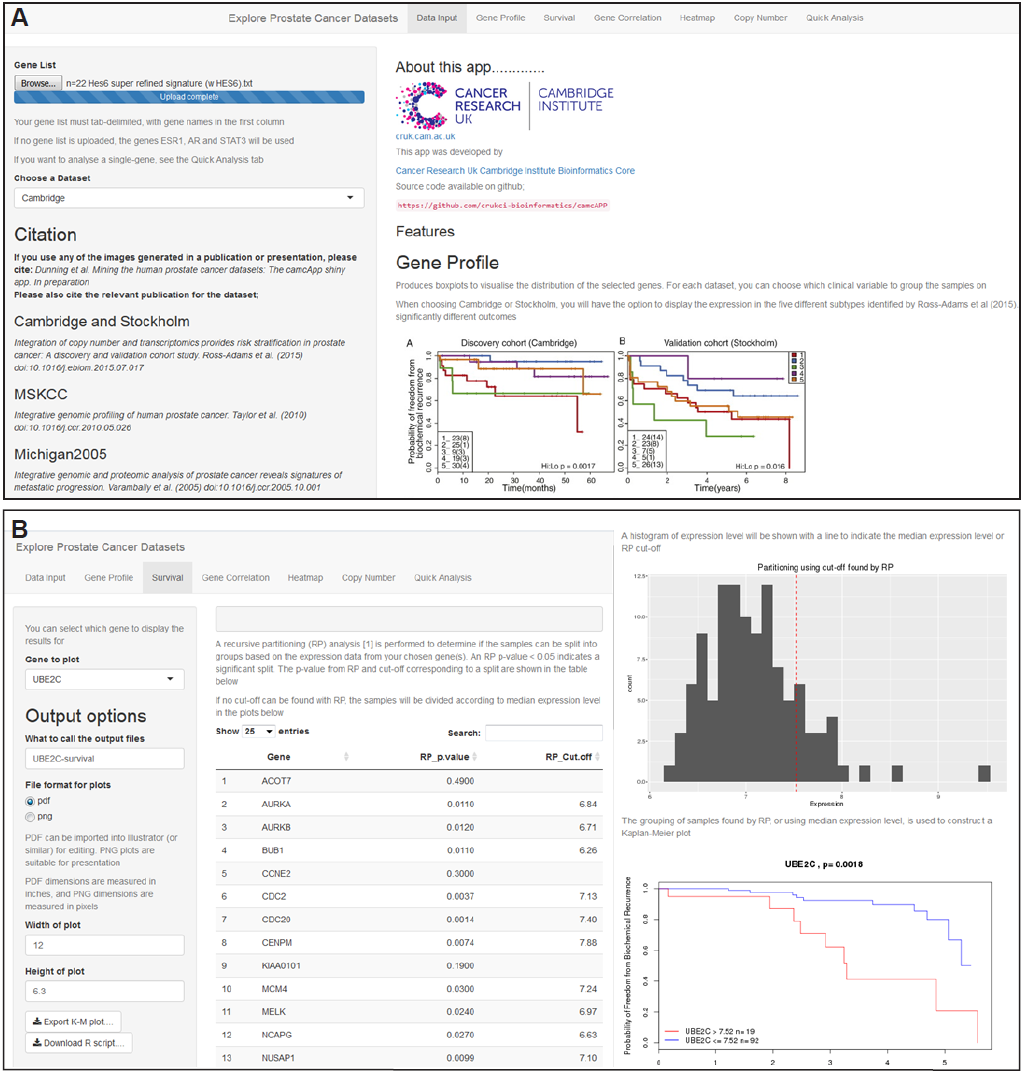
Examples of “camcaPP” shiny app functions – Gene-list input and Survival analysis. **A.** Entry point to site including upload point for gene lists and citation information. **B.** Kaplan-Meier biochemical relapse-free survival plots can be created for any selected gene from the input list in any selected dataset. Example of .pdf output modified in Adobe Illustrator™ demonstrated in panel **A** (Ross-Adams et al).^1^

1. Creation of boxplots to visualise the distribution of expression values of each gene for different clinical covariates of interest (**Figure 2**).
2. A survival analysis to assess whether the expression level of each gene can predict relapse (**Figure 1**). The “party” R package^14^ is first used to see if the expression level can be split into two distinct groups. If such a separation can be found, then a Kaplan-Meier curve is generated from the survival times of samples in the different groups. If no significant separation of samples is found, the median expression level is used to define groups of samples with high or low expression.
3. Pairwise correlations of the expression level of all specified genes.
4. Construction of a heatmap to assess whether the chosen genes can split the dataset of interest into subgroups. Various methods of clustering and visualisation are supported.
5. Tabulating the number of copy-number amplifications and deletions observed in the Cambridge, Stockholm and MSKCC cohorts, and making a heatmap of copy-number calls (Figure 2).^15^

**Figure 2.**
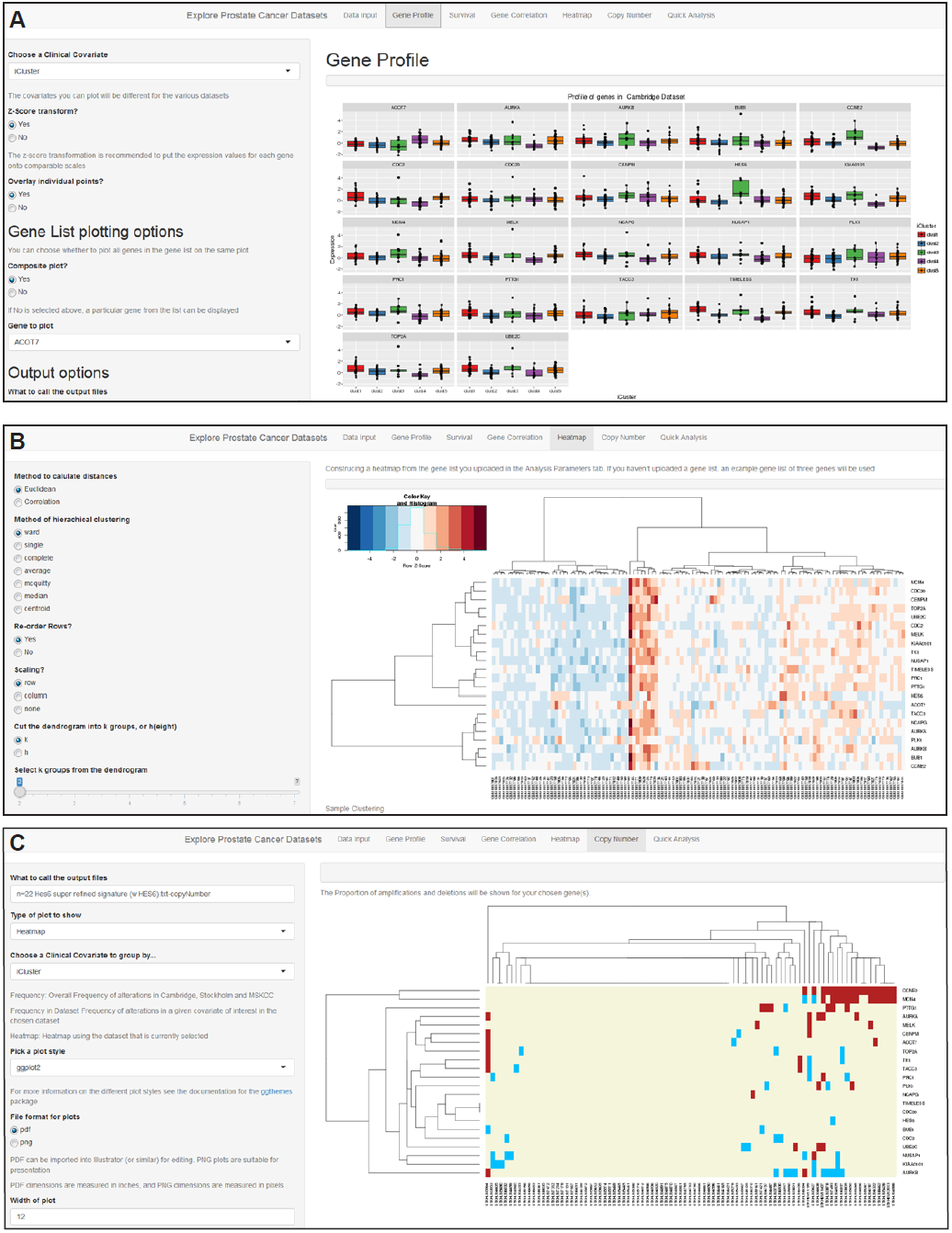
Examples of “camcaPP” shiny app functions – Gene expression and Copy Number plots. **A.** Boxplots for gene expression can be created for a list of genes – in this example the five Cambridge subgroups are demonstrated for a number of input genes. **B.** Heatmaps for gene expression can be created from any of the datasets. **C.** Copy number (CN) plots depicting CN gain or loss (or no change = neutral) can also be created. **D.** “Quick Analysis” tab allows rapid spot checks for single genes of interest focussing on relative expression, copy number and survival curves (example shown here is a boxplot showing relative gene expression in CRPC, tumour and benign tissue for HES6, a known driver of castration resistance^17^)

One of the challenges in constructing such a tool is delivering an output format that is readily transferable to slides for presentations or panels of a figure for publication. We recognise that this is, in part, a matter of axis typesetting and plot configuration but also of delivering an output file which permits further adjustment of the figure in, for example, Adobe Illustrator™. Thus all plots can be exported as PDF or PNG files with configurable dimensions. Furthermore, for those that are well-versed in R, the code to produce a particular plot can be downloaded and modified as required. A further challenge that we seek to address with this interface is merging datasets for combined analysis. We hope to offer this option in due course, as we include further datasets that include samples analysed on compatible platforms.

Strategies to address the Big Data problem have focussed on making the ever-increasing volume of genomic data accessible to scientists and on opening up the possibility of engaging nonspecialists.^16^ This approach embodies a responsible attitude to science both in terms of patient input and financial resource and we believe that tools such as this are an important step to maximising the value of these landmark studies.

